# Not quite human, not quite machine: Electrophysiological responses to robot faces

**DOI:** 10.1101/2020.06.11.145979

**Authors:** Allie R. Geiger, Benjamin Balas

## Abstract

Face recognition is supported by selective neural mechanisms that are sensitive to various aspects of facial appearance. These include ERP components like the P100, N170, and P200 which exhibit different patterns of selectivity for various aspects of facial appearance. Examining the boundary between faces and non-faces using these responses is one way to develop a more robust understanding of the representation of faces in visual cortex and determine what critical properties an image must possess to be considered face-like. Here, we probe this boundary by examining how face-sensitive ERP components respond to robot faces. Robot faces are an interesting stimulus class because they can differ markedly from human faces in terms of shape, surface properties, and the configuration of facial features, but are also interpreted as social agents in a range of settings. In two experiments, we examined how the P100 and N170 responded to human faces, robot faces, and non-face objects (clocks). We found that robot faces elicit intermediate responses from face-sensitive components relative to non-face objects and both real and artificial human faces (Exp. 1), and also that the face inversion effect was only partly evident in robot faces (Exp. 2). We conclude that robot faces are an intermediate stimulus class that offers insight into the perceptual and cognitive factors that affect how social agents are identified and categorized.

## Introduction

Face recognition is supported by neural mechanisms that are selective for different aspects of face structure. For example, face-sensitive neural loci that have been identified by fMRI studies include the OFA (Pitcher et al., 2007), the FFA (Kanwisher & Yovel, 2006) and the STS (Schohbert et al., 2018), which are part of an extended “face network” in extrastriate visual cortex (Wang et al., 2016). These different cortical regions each have different profiles of sensitivity and selectivity that suggest different functional roles in processing face stimuli. The occipital face area appears to implement some form of part-based processing that contributes to local feature analysis within a face pattern (Pitcher, Walsh & Duchaine, 2011). By contrast, the fusiform face area (FFA) appears to be tuned to larger-scale, perhaps “holistic,” aspects of facial structure (Zhang et al., 2012). Finally, the superior temporal sulcus exhibits sensitivity to multiple socially-relevant aspects of face appearance.(Deen et al., 2015). Analogous results from EEG and ERP studies of face recognition further support these functional divisions. The P100 ERP component, for example, shares some of the part-based selectivity of the OFA (Hermann et al., 2005), though it also exhibits effects of face inversion more commonly associated with holistic face processing (Colombatto & McCarthy, 2017). The N170 ERP component, certainly the most widely studied face-sensitive ERP component, exhibits a profile of sensitivity and selectivity that suggests it functions more or less as a face detector that relies on large-scale face appearance. For example, isolated face parts give rise to weaker N170 responses than full face patterns in some circumstances (Bentin et al, 1996), though the N170 also exhibits sensitivity to the appearance of the eyes in particular (Itier et al., 2007). Refining our understanding of the image properties that elicit strong responses from these various face-specific neural loci is an important step towards characterizing neural pathways for face recognition more completely. Further investigation of the boundary between “face” and “non-face” relative to specific neural responses is a powerful means of defining the underlying representation of face appearance that supports face processing at each stage in terms of both low-level image features and high-level category information.

A useful way to examine the boundary between faces and non-faces at specific stages of neural processing is to identify stimulus categories that are ambiguous with regard to their status as faces. Such stimuli may either possess or lack critical visual features that define the scope of face processing with regard to a specific neural response, thereby informing our understanding of the underlying representation of face appearance via the response to the ambiguous category. For example, several studies have examined aspects of pareidolia, which refers to the perception of faces in patterns that do not truly depict faces (e.g. faces in the clouds). Pareidolic stimuli are objectively not faces, but tend to share critically important features with face patterns (Omer et al., 2019), leading observers to label non-faces as faces under some circumstances (Paras & Webster, 2013), dwell longer on pareidolic non-faces than other non-face patterns, and make websites with amusing pictures of home appliances that look like they have a personality (see the Faces in Things account (@FacesPics) on Twitter). In terms of neural responses, pareidolic faces elicit more activity from the FFA as a function of how face-like they appear to naive observers (Wardle et al., 2017). Non-face patterns that triggered false alarms from a computational model of face detection (a sort of AI-defined pareidolia) also modulate N170 and FFA responses depending on how face-like they appear (Moulson et al., 2011; Meng et al., 2012; Nehi et al., 2018). Besides simply providing an interesting and whimsical stimulus category, these investigations of pareidolic faces offer a useful look at the boundary between faces and non-faces in extrastriate cortex. By measuring neural responses to these stimuli that are not quite faces, but not quite *not* faces, specific links between image features and neural responses have been established that suggest properties of the underlying representations maintained at different stages of face processing. For example, Paras & Webster (2013) found that while placing symmetric eye spots in 1/f “totem pole” images elicited pareidolia, this was not typically sufficient to elicit a robust N170 response. This kind of demonstration of what does and does not allow an image to cross the boundary from face to non-face offers useful information about the shape and position of that boundary, and therefore the nature of face tuning.

In the current study, we chose to investigate the boundary between face and non-face processing using robot faces, which we suggest are another example of a stimulus category that is useful for characterizing the neural basis of face processing due to its ambiguous status. Robot faces are a particularly interesting boundary condition to explore, as they differ from humans regarding surface, shape, and material properties, but can still be social beings. Though the appearance of robot faces differs from humans, their facial organization and pose can affect human-robot interactions (Ghazali, Ham, Barakova & Markopoulos, 2018; Broadbent et al., 2013), much like human expressions. Robots’ resemblance to human faces is also sufficient for infants to expect adults to speak with robots in their environment (Arita et al., 2005), though they do not exhibit such expectations regarding other objects. The degree of resemblance between human and robot faces is of critical importance, however: robot face likability decreases as robot appearance approaches human-like appearance, only rebounding when robot appearance becomes nearly human (Mathur & Reiching, 2016). Robot face displays that look humanoid are also rated as more likely to have a mind and to be alive than robot faces that have metallic appearance (Broadbent et al., 2013). Together, these results suggest that robot faces have the potential to be endorsed as social agents, though the strength of this endorsement may depend on a range of perceptual factors that lead robot faces to be either included or excluded from categorization as a face. We suggest that by examining the neural response to robot faces, we will gain further insight into how face-specific mechanisms may or may not be engaged by faces that differ substantially from the faces observers typically interact with, but nonetheless can be understood as social agents with mental states and intentions.

Our present goals were two-fold: (1) To compare the N170 response elicited by robot faces to human faces and an unambiguous non-face category, (2) To compare the N170 face inversion effect (FIE) across human face, robot face, and non-face categories. In both cases, we use the response properties of the N170 as a way to measure how face-like the neural response to robot faces is relative to human faces and objects. In Experiment 1, we used the amplitude and latency of the N170 as proxies for face-ness in upright images, while in Experiment 2, we used the difference in response between upright and inverted faces as another proxy for how face-like each category is according to the N170. Briefly, we found in both experiments that robot faces occupy a middle ground between faces and non-faces. In terms of the N170, robot faces elicit an intermediate response that suggests they tend to share some critical features with human faces, but also lack some important image structure that the N170 is tuned to. We also find differential effects at the P100 and N170 that suggest that the status of robot faces as faces may vary depending on what stage of processing we consider. Specifically, robots’ substantial deviation from human-like appearance in local neighborhoods may have larger consequences at earlier stages of face processing, while their general adherence to the first-order geometry of human faces may lead them to be processed in a more face-like way at later stages of processing that entail holistic processing. We close by discussing the potential for more targeted analysis of critical image features that may lead robot faces to look more-or-less face-like, and the possibility that social interactions between robots and humans may affect the status of robot faces within the extended face network.

### Experiment 1 - How do face-sensitive ERP components respond to robot faces?

In our first experiment, our goal was to compare the response of face-sensitive ERP components (the P100 and N170) to human faces, robot faces, and non-face objects.

## Methods

### Participants

Participants between the ages of 18-22 years old were recruited from the NDSU Undergraduate Psychology Study Pool. Our final sample was composed of 24 participants (15 female) who self-reported normal or corrected-to-normal vision and most of which were right-handed (n=2 with mixed left-handed responses), as assessed by the Edinburgh Handedness Inventory (Oldfield, 1971). Prior to the beginning of the experiment, we obtained informed consent from all participants.

### Stimuli

The doll, robot, and clock stimuli in this study were obtained from Google Image searches for those object categories, while photographic faces and their computer-generated (CG) counterparts were selected from a pre-existing database maintained by our laboratory. Thirty images of each stimulus were included in our final task. We created these stimuli by converting all images greyscale and resizing them to 512×512 pixels. We then used the MATLAB SHINE Toolbox (Willenbockel et al., 2011) to equalize the power spectra of all stimulus images (Figure 1), to minimize the possibility that any effects we observed at either target component were the result of category differences in low-level image properties.

**Figure 1.**
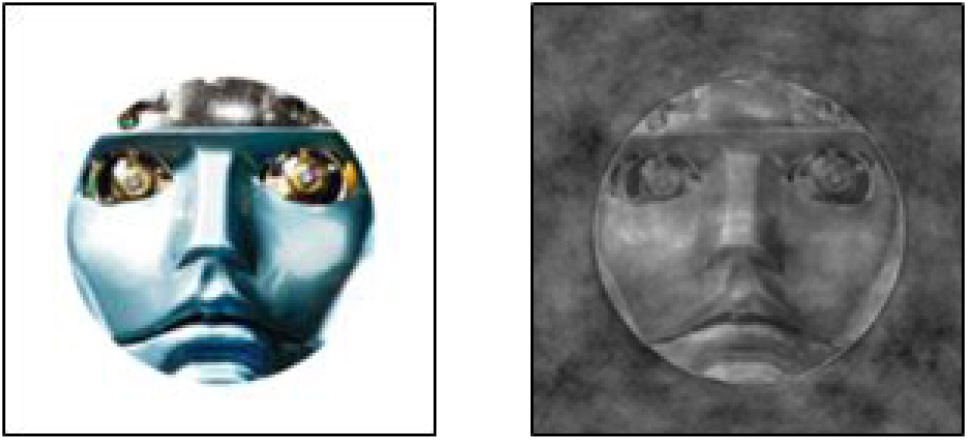
The left image represents an unedited robot face image. The image on the right is an example of the same image following the application of power spectrum matching implemented using the MATLAB SHINE Toolbox.

### Procedure

After obtaining consent, we measured the circumference of each participant’s head to ensure the selection of a properly fitting Hydrocel Geodesic Sensor Net by EGI. Each net was soaked in a potassium chloride (KCl) solution for 5 minutes before being placed on the participant’s scalp. The net was then plugged into an EGI 400 NetAmps amplifier. Following, the impedance of each electrode was measured and those above 25kW were adjusted and/or applied with more KCl solution until an adequate measurement was met.

The EEG recordings took place in an electrically-shielded and sound-attenuated chamber where our participants sat in front of a 1024×768 LCD monitor. EEG data was collected at a sampling rate of 250 Hz and NetStation v5.0 was used for the EEG recordings and event markings. Participants were presented with 30 unique images of each stimulus category. Each image was randomly presented twice in a pseudo-randomized order determined by functions in EPrime v2.0 during the session for a total of 300 images. The images were displayed on a white background for 500 milliseconds with random interstimulus intervals (ISI) between 800 and 1500 milliseconds. We asked participants to respond to these images by pressing a ‘person-like’ button for images of human, computer-generated, and doll faces, with one of their hands, and a ‘machine-like’ button for clock and robot faces with the other. To account for the expected right-hand preference, the orientation of the response-box was turned 180 degrees after every participant, so that half of participants used their right hand for the ‘person-like’ button, and the other half used their left. This task took about 15 minutes and the behavioral data was collected using EPrime v2.0.

## Results

### Behavioral Data - Human/Machine Classification

For each participant, we calculated the median response time for correct human/machine judgments in each category. For all images of human faces, the correct answer was “human,” while robot faces and clocks should have been labeled “machine.” We analyzed these values using a 1-way repeated-measures ANOVA implemented in JASP (JASP Team, 2018) with image category (real faces, CG faces, doll faces, robot faces, and clocks) as a within-subjects factor. This analysis revealed a significant main effect of image category (F(4,108)=4.06, p=0.004), and post-hoc testing revealed a significant pairwise difference between robot faces and clocks such that response latencies to robot faces were significantly slower (Mean diff. = 27.0ms, s.e. = 8.08ms, t=3.35, p_bonf_=0.024). Average response latencies across participants are displayed in Figure 2.

**Figure 2.**
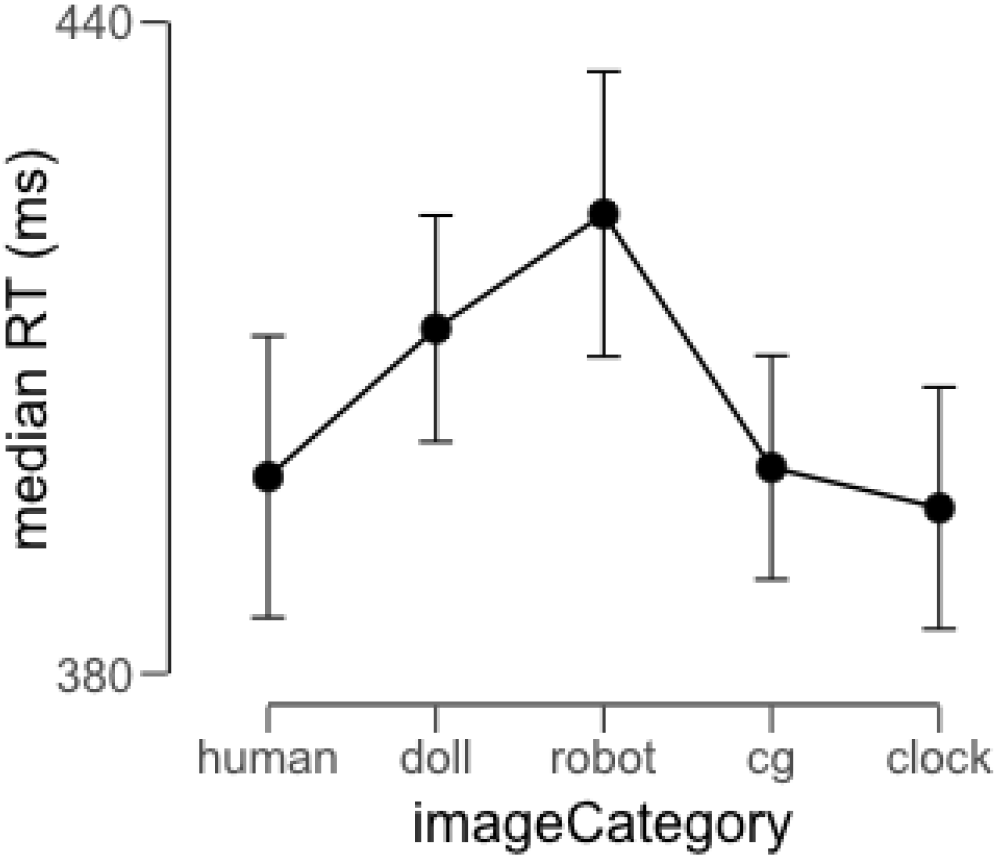
Average response latency for correct human/machine judgments across image categories. Error bars represent 95% credible intervals.

### ERP component analyses

We continued by examining both the mean amplitude and the peak latency of the P100 and N170 ERP components elicited by stimuli from each image category. For each participant, the continuous EEG recorded during the testing session was filtered using a 0.1Hz-30Hz bandpass filter, and then segmented into 1100ms epochs using markers inserted into the EEG record during data acquisition. Each of these segments began 100ms before stimulus onset and ended 1000ms after stimulus onset. Each segment was baseline corrected by subtracting the average value calculated during the 100ms baseline period from the entire waveform. Subsequently, we applied algorithms for identifying and removing eye movement artifacts from the data and used spherical spline interpolation to remove and replace any systematically bad channels in the sensor array. Finally, we calculated average ERPs for each participant by averaging together the data within each image category at each sensor, followed by the application of an average re-reference.

We identified time intervals and sensors of interest by visually inspecting a grand average ERP calculated across participants (Figure 3), and locating maximal deflections corresponding to the components of interest. We analyzed both components using three sensors in the left hemisphere (sensors 29, 32, and 33) and three sensors in the right hemisphere (43, 44, and 47). We selected an interval of 92 ms-136 ms to measure P100 amplitude and latency, and an interval of 140 ms-192 ms to measure N170 amplitude and latency. For each ERP descriptor, we calculated these values for each participant and analyzed the resulting data using a 5×2 repeated measures ANOVA with image category (human face, doll face, cg face, robot face, and clock) and hemisphere (left or right) as within-subjects factors.

**Figure 3.**
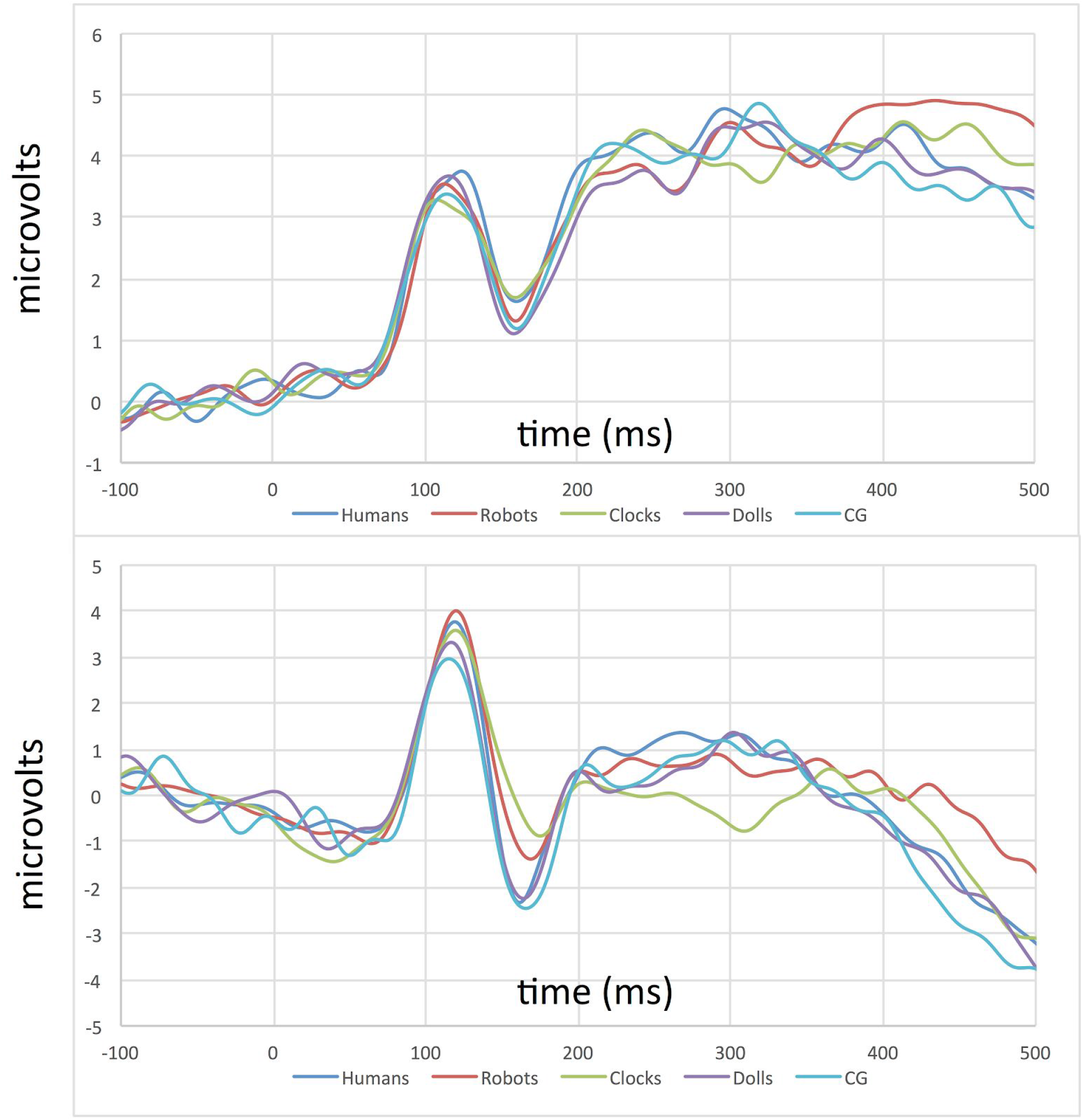
Grand average ERPs from the left (top) and right (bottom) hemispheres depicting the P100/N170 complex for all stimulus conditions in Experiment 1.

### P100 Amplitude

Our analysis of the P100 mean amplitude yielded neither a significant main effect of image category (F(4,92)=1.56, p=0.19) nor a main effect of hemisphere (F(1,23)=2.06, p=0.16). The interaction between these two factors also did not reach significance (F(4,92)=1.61, p=0.18).

### P100 Latency

Our analysis of the P100 latency-to-peak data yielded a significant main effect of image category (F(4,92)=2.86, p=0.028). In subsequent post-hoc tests, no pairwise difference between categories reached significance following corrections for multiple comparisons, but pairwise differences between human and CG faces (t=−2.77, p=0.079), doll faces and CG faces (t=−2.76, p=.083) and doll faces and clocks (t=−2.69, p=0.098) were marginally significant (Figure 4). Neither the main effect of hemisphere (F1,23)=0.18, p=0.68) nor the interaction between these two factors (F(4,92)=1.48, p=0.22) reached significance.

**Figure 4.**
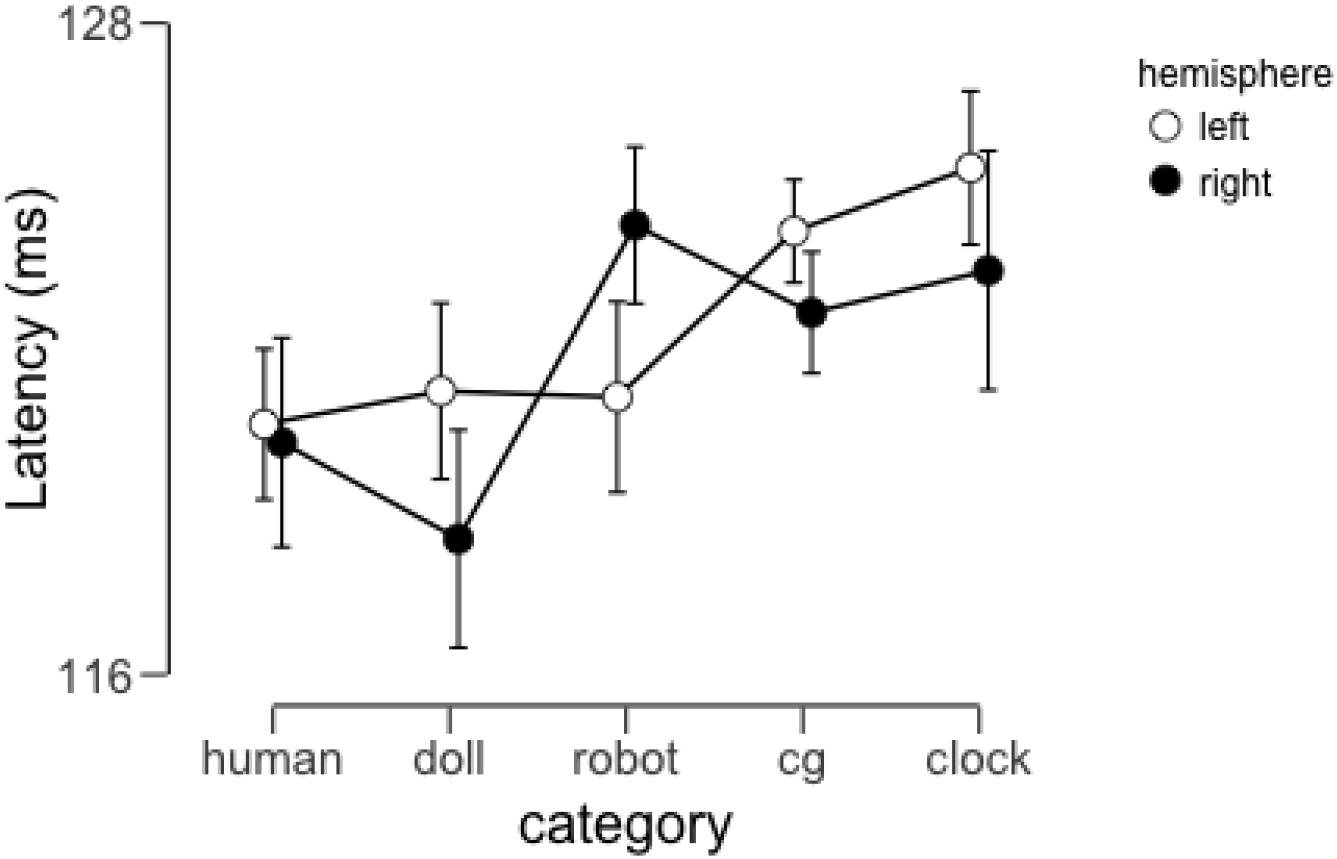
Average P100 peak latencies across participants as a function of image category and hemisphere. Error bars represent 95% credible intervals.

### N170 Amplitude

This analysis revealed a main effect of image category (F(4,92)=2.85, p=0.028), but neither a main effect of hemisphere (F(1,23)=2.76, p=0.11) nor a significant interaction between these factors (F(4,92)=1.38, p=0.25). Post-hoc tests revealed that the main effect of image category was the result of significant pairwise differences between robot faces and CG faces (t=3.34, p=0.016) and marginal pairwise differences between human and robot faces (t=2.87, p=0.06) and between doll faces and robot faces (t=2.99, p=0.057). In Figure 5 we display average mean amplitude across participants as a function of image category.

**Figure 5.**
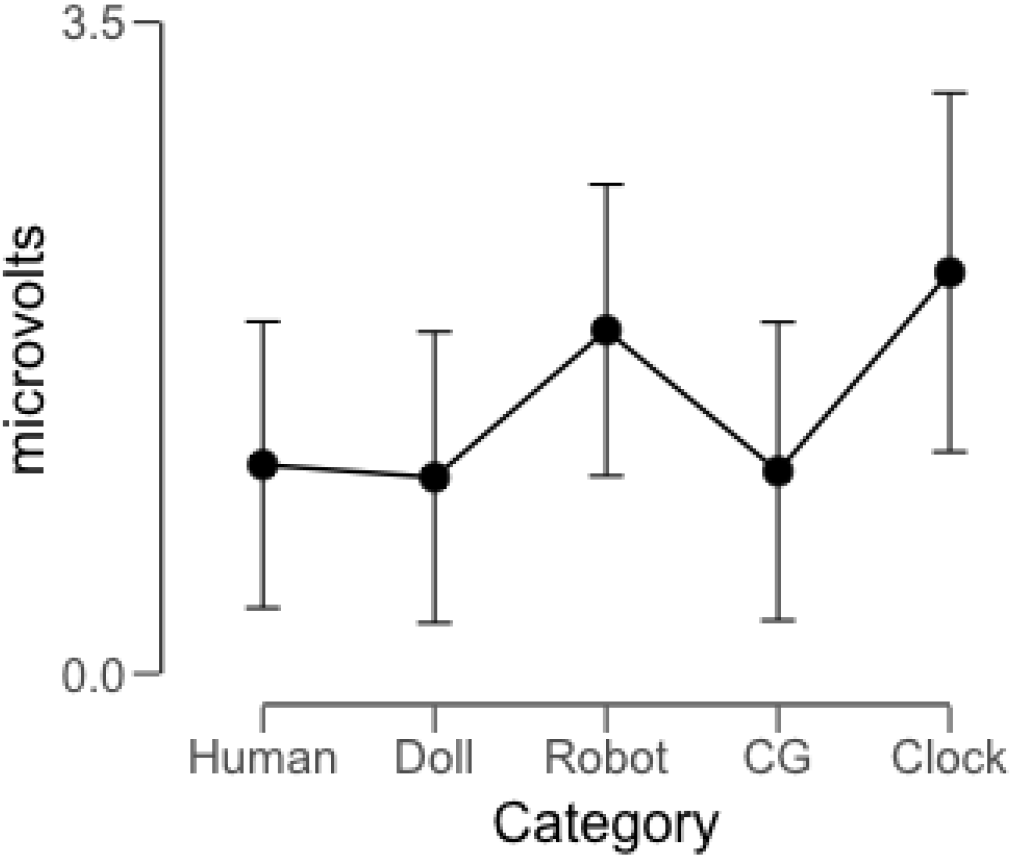
Average N170 mean amplitude across participants as a function of image category (collapsed across hemisphere). Error bars represent 95% credible intervals.

### N170 Latency

This analysis revealed a significant main effect of image category (F(4,92)=4.48, p=0.002), but neither a main effect of hemisphere (F(1,23)=0.36, p=0.55) nor an interaction between these factors (F(4,92)=1.88, p=0.12). Post-hoc tests revealed that the main effect of image category was the result of pairwise differences between human faces and clocks (t=−3.63, p=0.007), doll faces and clocks (t=−3.07, p=0.035), and robot faces and clocks (t=−4.43, p<0.001). In Figure 6, we display the average N170 latency across participants as a function of image category and hemisphere.

**Figure 6.**
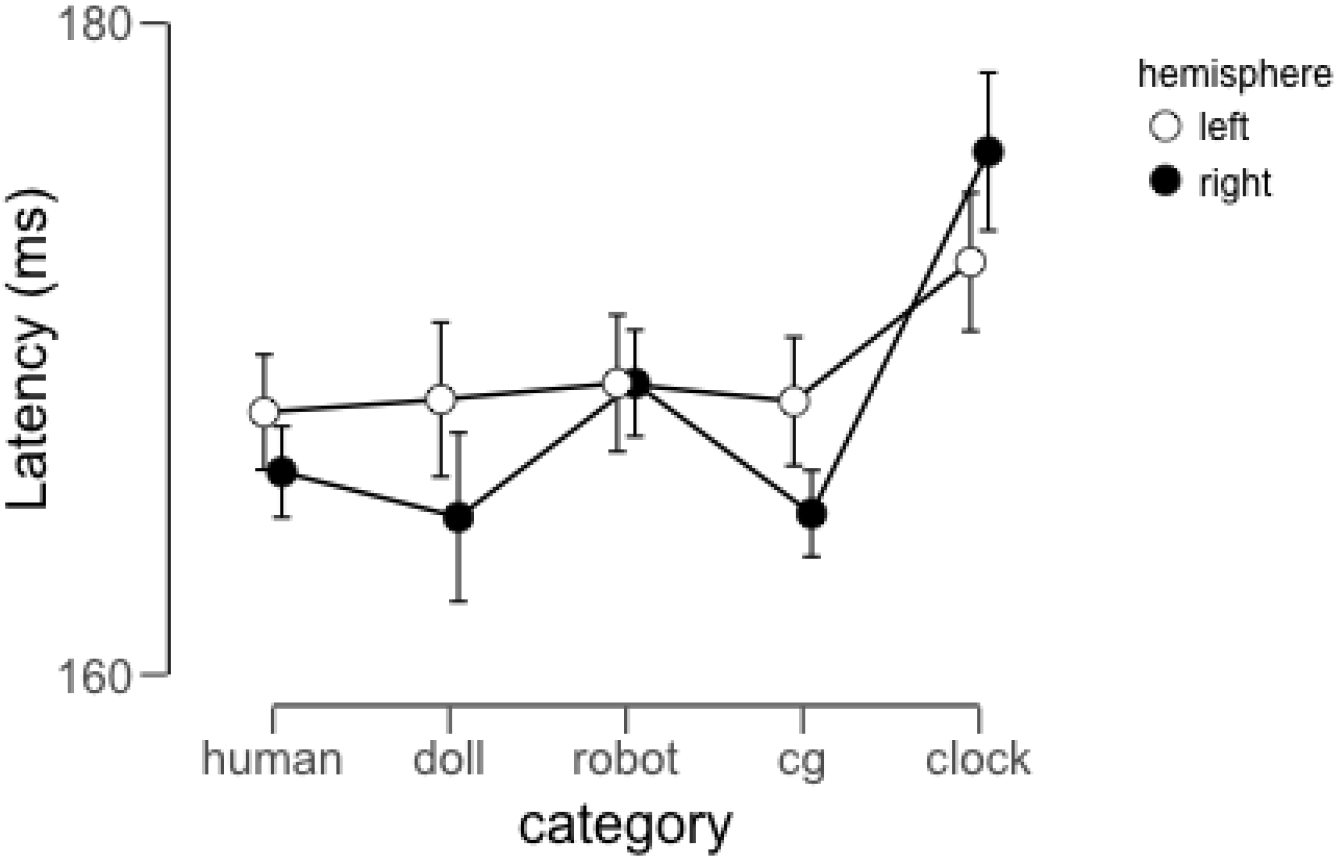
Average N170 Latency across participants as a function of image category and hemisphere. Error bars represent 95% credible intervals.

## Discussion

The results of our first experiment provide converging evidence from different descriptors of the ERP waveform that robot faces lie on the boundary between faces and non-faces, while artificial human faces elicit responses that are decidedly face-like. This latter feature of our data is consistent with previous reports indicating limited effects of artificial appearance on the N170 amplitude and latency (Balas & Koldewyn, 2013; Wheatley et al., 2011). There are substantial differences in appearance between real human faces, dolls, and computer-generated faces, and artificial appearance is known to disrupt performance in a range of face recognition tasks. Artificial faces are harder to remember, for example (Balas & Pacella, 2015), and elicit different social evaluations of qualities like trustworthiness even when identity is matched between real and artificial images (Balas & Pacella, 2017). Despite the existence of a perceptual category boundary between real and artificial faces (Looser & Wheatley, 2010) and the various recognition deficits that appear to accompany artificial face appearance, the neural processing of these images at the N170 is similar to that of real human faces. This is an important observation because it demonstrates that the tuning of the N170 to face appearance is broad enough to include these categories, and also because it suggests that strong attribution of a mind, intentions, or other dimensions of social agency (which are diminished for artificial faces, Balas & Pacella, 2017) are not necessary to elicit a robust N170 response.

Compared to these artificial human face conditions, the differences between our robot condition and human faces are more complex. While the profound differences in facial appearance between these categories did not elicit differential processing at the P100, a complicated picture emerges from the N170’s response properties. The amplitude data suggests weak differentiation of robot faces from human faces, and no evidence of a difference between robots and clocks. We suggest that these outcomes support the hypothesis that robots are perceived more like objects than faces. However, the N170 latency data reveals that robot faces (along with other human face categories) differ from non-face objects, suggesting a more face-like response with regard to this descriptor. We propose that these somewhat conflicting results reflect the intermediate status of robot faces - neither entirely face-like, nor entirely machine-like, but straddling the boundary between faces and non-faces in terms of these neural responses.

While this is an interesting result, our first experiment does not allow us to rule out an important alternative account, however. Specifically, the amount of variability across human faces, robot faces, and clocks was not explicitly measured or controlled in our stimulus set, and any differences in within-category image variability could contribute to the results we observed. Inter-stimulus perceptual variance (ISPV) has been shown to modulate N170 amplitudes in particular (Thierry et al., 2007), with larger ISPV values associated with smaller N170 amplitudes due to increased trial-by-trial jitter of the N170. Our results are therefore consistent with a stimulus set that has increased ISPV for robot faces compared to human faces. This could be the case in our stimulus set; Robot faces vary substantially across manufacturers according to the roles they are intended to occupy, while our CG faces were matched to the identities of our real human faces. The intermediate status of robot faces could thus reflect an artifact of our stimulus set rather than the ambiguous category status of robots.

To help address this issue and further examine the extent to which face-specific processing includes robot faces, in our second experiment we chose to investigate the face inversion effect (FIE) at our target components for human faces, robot faces, and clocks. In terms of ERP responses, the FIE refers to the increased amplitude and latency of the N170 component in response to inverted face images (Rossion et al., 2000). We suggest that this is a useful way to address the issue of potential category differences in ISPV because the effect of inversion on amplitude follows from an ISPV-based account only if we assume that inversion increases stimulus homogeneity, which is itself an indicator of orientation-dependent processing. Further, latency effects on the N170 with inversion are to our knowledge not easily explained in an ISPV account (Rossion & Jacques, 2008), making these markers of the FIE a more robust indicator of face-specific processing for a particular stimulus category. We hypothesized that if robot faces are indeed a boundary category that lies between faces and non-faces, we may see an FIE that is intermediate to that observed for human faces and clocks. Alternatively, if the FIE for robot faces clearly mirrors what we observe in either comparison category, this may make a stronger case for robots being considered unambiguously as either faces or non-faces at these stages of visual processing.

### Experiment 2 - Do robot faces yield a face inversion effect at face-sensitive ERP components?

In Experiment 2, we investigated the P100 and N170 responses to upright and inverted human, robot, and clock faces. Both components have been reported to exhibit a face inversion effect (FIE) such that inverted faces lead to larger amplitudes, and our goal was to use this effect as a proxy for face-like processing.

## Methods

### Participants

Our final sample was composed of 24 students in the NDSU Undergraduate Psychology Study Pool (Female=16) between the ages of 18-23. Only 1 participant reported mixed left-handedness (Oldfield, 1971) and all participants self-reported normal or corrected-to-normal vision. We obtained informed consent from all participants prior to the beginning of the experiment.

### Stimuli

For this experiment, we presented participants with right-side-up and inverted clock, robot, and human face images (Figure 7). Because the ERP components of doll and computer-generated facial stimuli were so similar to that of human face images in experiment 1, we excluded them from this study. The images for this study were taken from experiment 1 and therefore, did not undergo editing. However, many robot face images looked similar when inverted, so we handpicked robot face images that looked noticeably inverted and ended up with 18 images. We then randomly chose 18 images from the other stimulus categories. These 54 images were duplicated and inverted, leaving us with 54 upright and 54 inverted stimuli.

**Figure 7.**
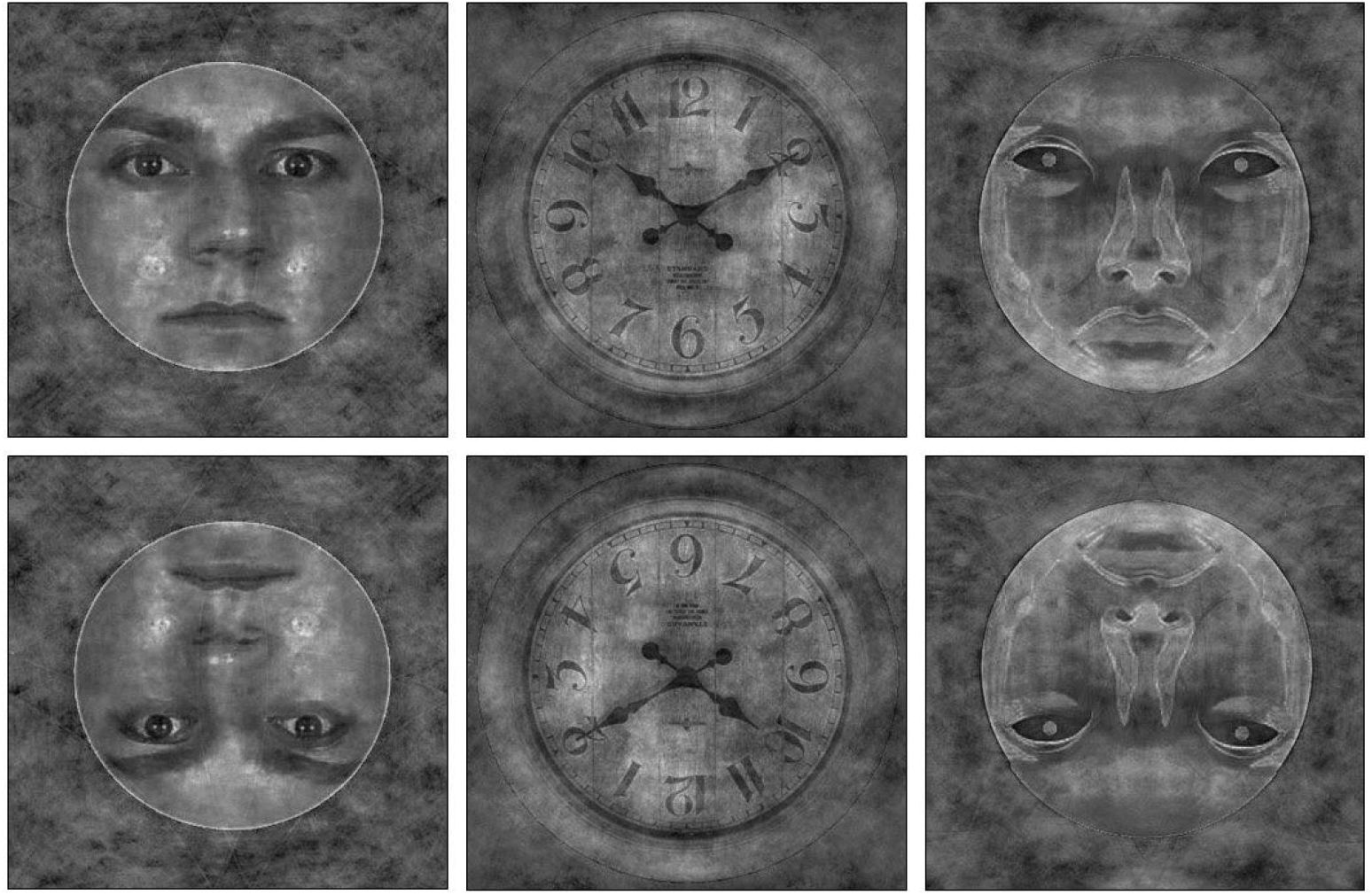
Example images from each stimulus category presented in upright and inverted orientations.

### Procedure

All recording software and procedures for preparing the participant and the Hydrocel Geodesic Sensor Net for the session were identical to those reported in Experiment 1.

Participants were presented with each stimulus in a pseudo-randomized order four times for a total of 432 trials. The images were displayed for 500ms on a white background with random ISIs ranging between 800 and 1500ms. Participants were asked to respond to the images by pressing a ‘person-like’ button for upright and inverted human faces and a ‘machine-like’ button for upright and inverted clock and robot face images. Participants were asked to use one hand for each button. The response box was flipped 180 degrees for half of our participants to balance which hand corresponded to which response cross participants.

## Results

### Behavioral Data - Human/Machine Classification

For each participant, we calculated the median response time for correct human/machine judgments in each category. For all images of human faces, the correct answer was “human,” while robot faces and clocks should have been labeled “machine.” We analyzed these values using a 2-way repeated-measures ANOVA implemented in JASP (JASP Team, 2018) with image category (real faces, robot faces, and clocks) and orientation (upright and inverted) as within-subjects factors. This analysis revealed only a marginally significant main effect of image category (F(2,44)=2.85, p=0.069). Neither the main effect of orientation (F(1,22)=1.67, p=0.21) nor the interaction between these factors (F(2,44)=0.80, p=0.46) reached significance. Post-hoc tests revealed that the marginal main effect of image category was the result of a marginally significant difference between response latencies to human and robot images (t=2.40, p=0.062), such that robot faces (M=456.35ms, s.d.=155.4ms) were classified correctly more slowly than human faces (M=438.1ms, s.d.=167.6ms).

### ERP component analyses

As in Experiment 1, we identified sensors and time windows of interest by inspecting the grand average ERP calculated across participants (Figure 8). We analyzed the P100 and N170 components at the same sensors as in Experiment 1, and used a time window of #ms-#ms to measure the P100 amplitude and latency, and a time window of #ms-#ms to measure the N170 amplitude and latency.

**Figure 8.**
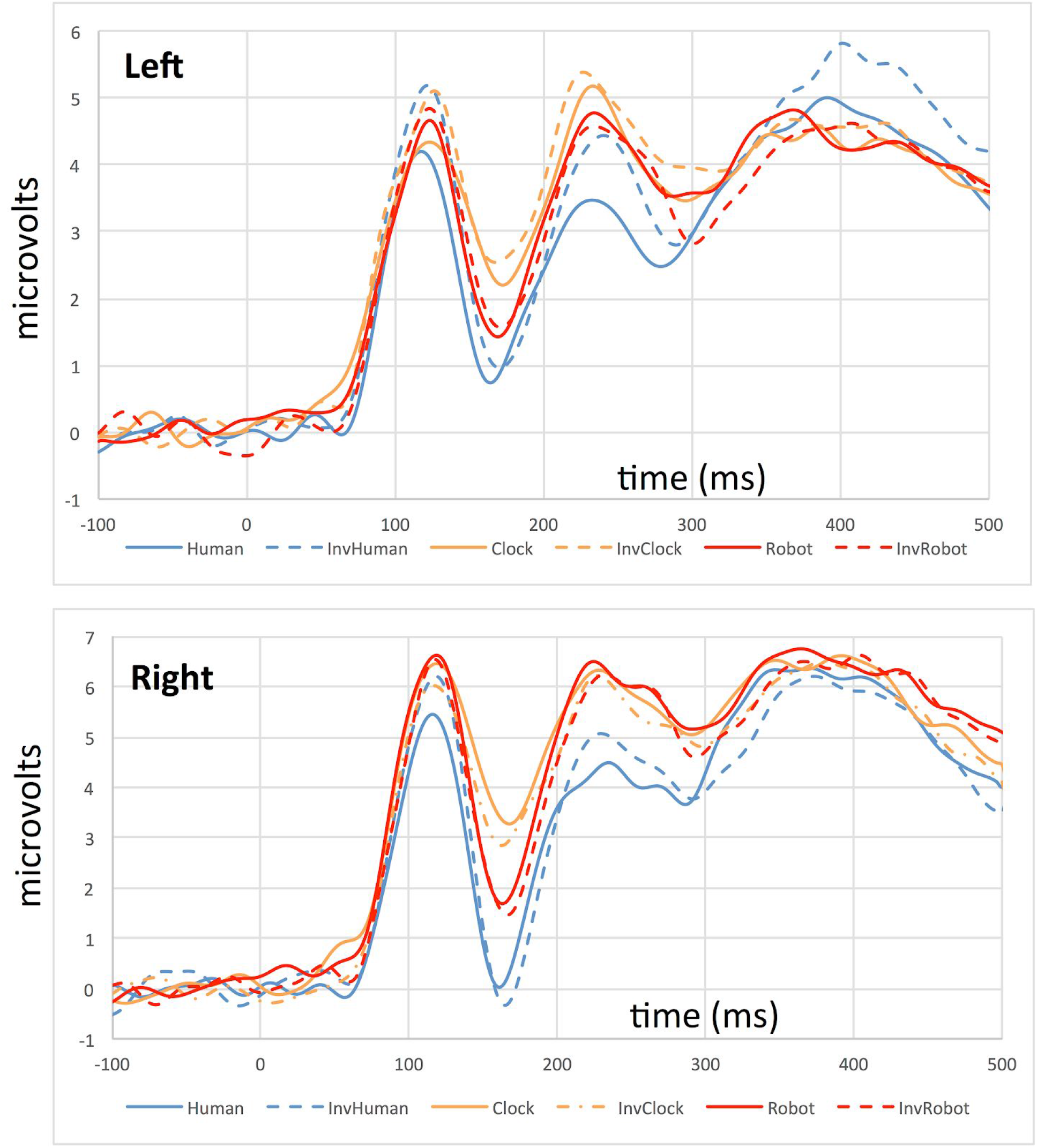
Grand average ERP waveforms for the left (top) and right (bottom) hemisphere depicting the P100/N170 complex for all conditions in Experiment 2. Solid lines depict the responses to upright stimuli, while dashed lines depict responses to inverted stimuli.

### P100 Amplitude

This analysis revealed a significant main effect of image category (F(2,44)=5.32, p=0.009), but no main effects of orientation (F(1,22)=2.10, p=0.16) or hemisphere (F(1,22)=3.22, p=0.086). We also observed significant interactions between image category and orientation (F(2,44)=14.7, p<0.001) and between orientation and hemisphere (F(1,22)=8.35, p=0.009). No other interactions reached significance

Post-hoc testing revealed that the main effect of image category was the result of a significant pairwise difference between responses to human faces and clocks (t=−3.84, p<0.001), and a marginally significant difference between robot faces and clocks (t=−2.23, p=0.084). The interaction between image category and orientation was the result of a significant orientation effect for human faces such that inverted faces elicited a more positive P100 peak than upright faces, but no such difference between upright and oriented clocks or robot faces (See Figure 9).

**Figure 9.**
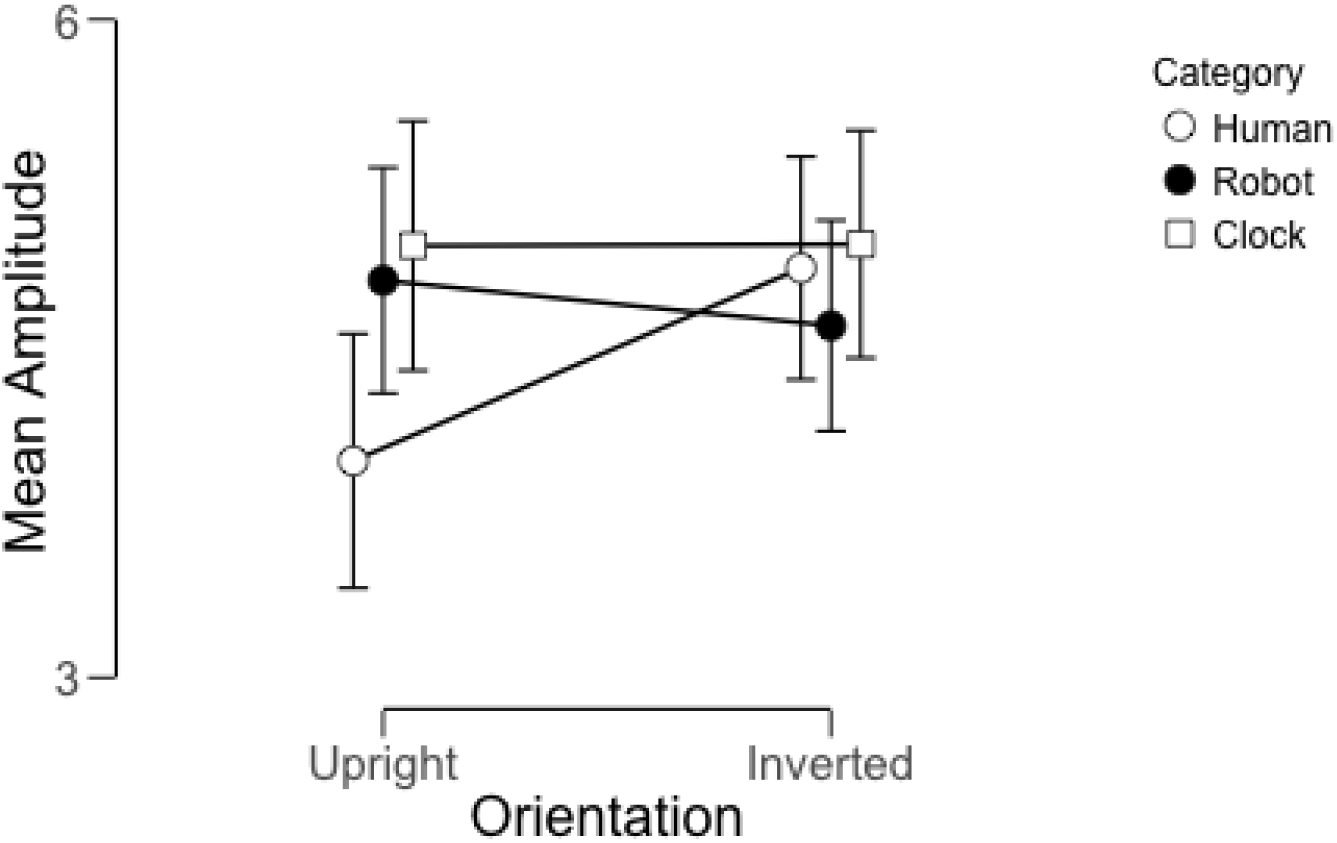
Average P100 amplitude across participants as a function of image category and orientation (collapsing across left/right hemisphere). Error bars represent 95% credible intervals.

### P100 Latency

We observed no significant main effects or interactions on the P100 latency to peak. Neither image category, orientation, or hemisphere affected this aspect of participants’ waveforms.

### N170 Amplitude

This analysis revealed a main effect of image category (F(2,44)=40.2, p<0.001), but no main effects of orientation (F(1,22)=0.55, p=0.41) or hemisphere (F(1,22)=1.54, p=0.23). The lack of a main effect of orientation is surprising, but we did also observe a significant interaction between orientation and hemisphere (F(1,22)=4.44, p=0.047). No other interactions reached significance.

The main effect of image category was driven by significant pairwise differences between all three image categories (t>6.11 in all cases, p<0.001 for all). The interaction between orientation and hemisphere was driven by a larger difference between left and right hemisphere responses for upright faces than for inverted faces (Fig. 10), which we interpret as a reflection of a typical N170 inversion effect in the right hemisphere, but not the left.

**Figure 10.**
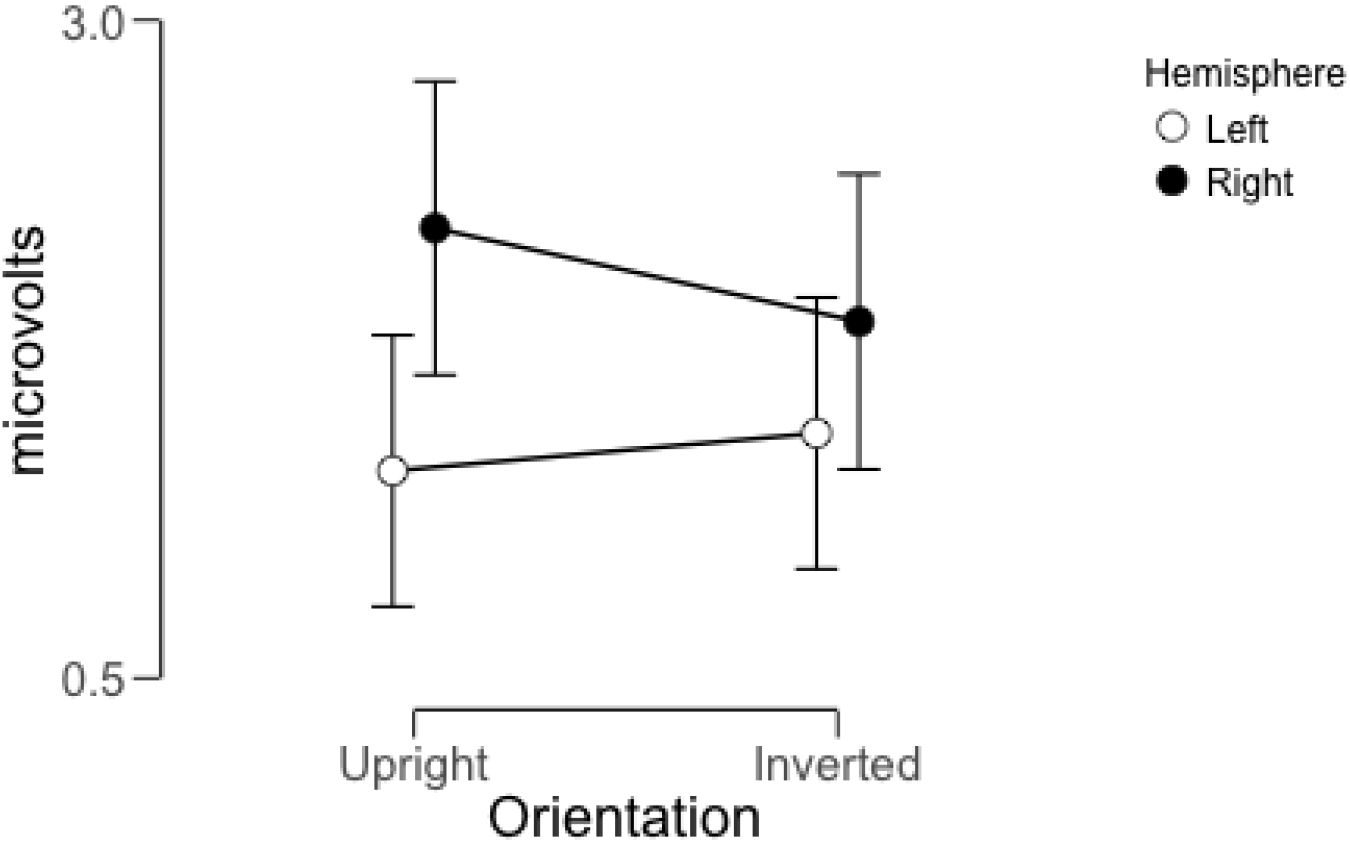
Average N170 amplitude across participants as a function of image orientation and hemisphere (collapsed across image category). Error bars represent 95% credible intervals.

### N170 Latency

This analysis revealed only a significant interaction between image category and orientation (F(2,44)=3.2, p=0.05). No other main effects of interactions reached significance. Post-hoc testing revealed that this interaction was driven by significant inversion effects for human and robot faces such that inverted faces elicited slower latencies to peak than upright faces, but no effect of image orientation on clock images (Figure 11).

**Figure 11.**
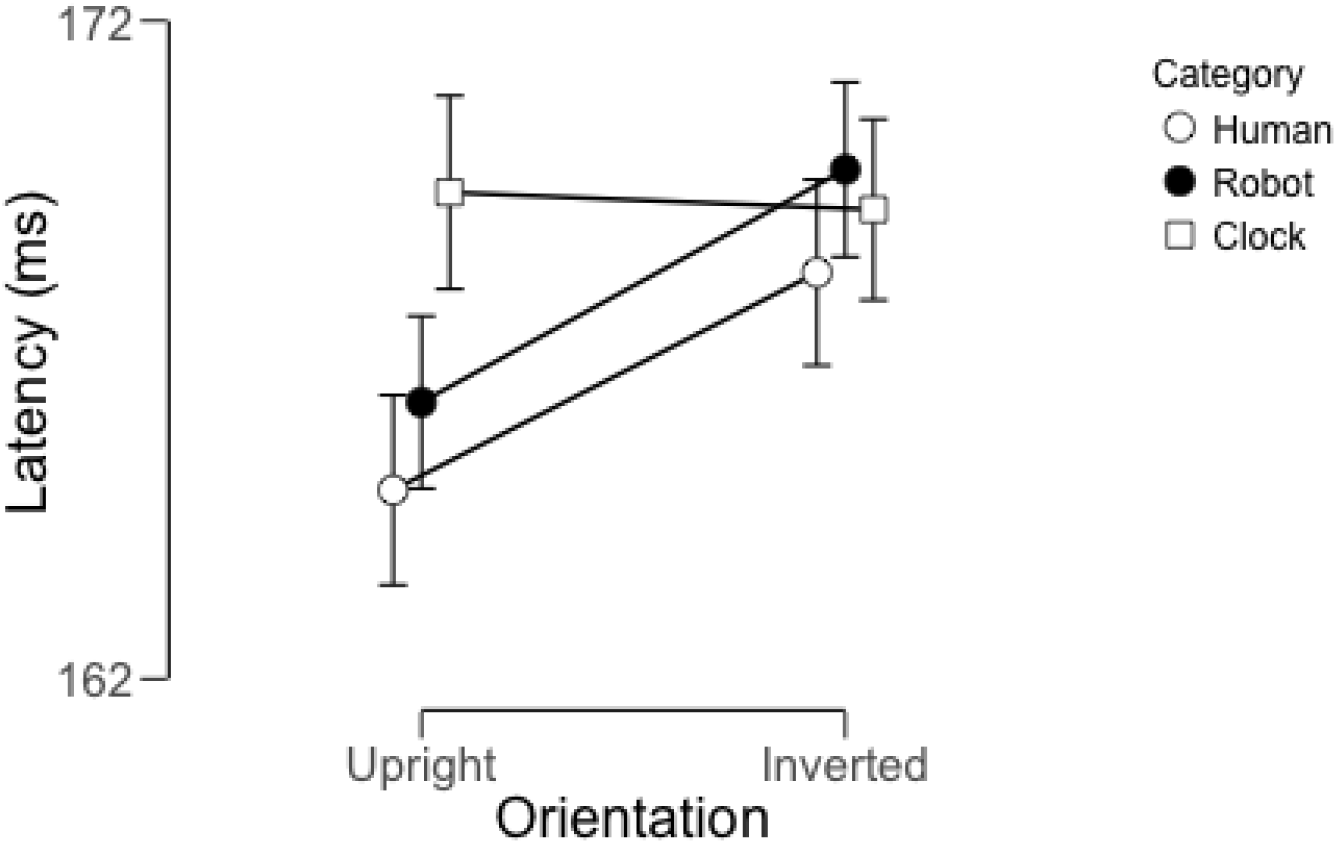
Average N170 latencies across participants as a function of image category and orientation. Error bars represent 95% credible intervals.

## General Discussion

Across two experiments, we have found that the neural response to robot faces has properties that suggest robot faces indeed lie on the boundary between faces and non-face objects. In our first experiment, we found that the N170 amplitude elicited by robot faces was intermediate to the response to human faces of multiple types and non-face objects. In Experiment 2, we further demonstrated that the face inversion effect was generally not evident for non-face objects, but was evident to a limited extent for robot faces. To the extent that we should accept the inversion effect as a proxy for the engagement of face-specific processing, these results together suggest that robot faces are capable of engaging these mechanisms to a limited degree while still different enough from human faces to elicit a different response.

In particular, in Experiment 2 we found that the face inversion effect was not evident for robot faces at the P100, but was observed in the N170 latency data. Despite the fact that the P100 generally reflects low-level processing that can be affected by basic image properties, this component also demonstrates some sensitivity to upright vs. inverted face images (Colombatto & McCarthy, 2017). However, in terms of the extended face network in extrastriate cortex, this early component also appears to reflect a different stage of processing than the subsequent N170 (Desjardins & Segalowitz, 2013). In particular, the P100 has been associated with part-based, as opposed to holistic, face processing (Pitcher et al., 2011), but this interpretation is somewhat at odds with theoretical accounts of face inversion as an index of holistic processing. For our purposes, we think that the presence of a robot face inversion effect at the N170 but not at the P100 reveals an interesting functional difference between robot and human faces regardless of the precise mechanistic differences between the processes indexed by these two components. Whatever processes these two components reflect, there is some property of robot faces that passes muster with the later component but is not sufficient to elicit face-specific processing from the P100.

One intriguing possibility is that social categorization may play an important role in determining how the processes indexed by these two components engage with robot faces versus human faces. In terms of behavioral responses, there is some evidence that out-group faces are less susceptible than in-group faces to manipulations that affect holistic processing (Michel et al., 2006), including the face inversion effect (Wiese, 2013). Neural responses also appear to be sensitive to social categories as well: Colombatto & McCarthy (2007) reported that the face inversion effect at the P100 was only evident for White faces viewed by White observers, and not evident for Black faces. This is in contrast to prior reports that demonstrated sensitivity to face orientation for White and Black faces at the N170 (Balas & Nelson, 2010), which suggests that these two components differ in part based on the extent to which social information modulates the nature of the neural response. We therefore suggest that one account of our data is that the social other-ness of robots is sufficient to exclude them from face-specific processing at an early cortical stage, but does not exclude them from subsequent face-specific processing at the N170. Robots are indeed subject to social categorization effects that affect how human observers interact with them. Eyssel & Kuchenbrandt (2012) demonstrated that robots can be favored as in-group or out-group members based solely on the name/label they are given without further interaction. This suggests that robots, though they are objects, not only act as social beings but may also be assigned to social categories in the same way as human faces. To our knowledge, the extent to which perceptual consequences of the other-race effect apply to robot faces has not been pursued. One potentially useful follow-up to the current experiments would be an examination of the strength of the other-race effect for robot faces (defined by perceptual discrimination abiities or memory biases) compared to human faces in relation to the neural effects we have reported here.

Another interesting way to examine the role of social categorization in affecting the inclusion or exclusion of robot faces from face-specific processing is to manipulate how robots are socially categorized through experience and exposure. Several results demonstrate that robots are not just perceptually a sort of borderline category for face processing, but they also occupy a boundary area in terms of people’s social inferences about their ability to have mental states, feelings, and personality. As such, even brief social interactions can lead to changes in how human observers evaluate robot agents. A social exchange between humans and robots can change beliefs and actions towards robots (Horstman et al., 2018). People can feel empathy towards a robot from simply hearing a story about it, and furthermore, this empathetic feeling makes participants hesitate when told to strike it (Darling, Nandy, & Breazeal, 2015). Social exchanges between robots and children have also been shown to change more complex evaluations regarding robots. For example, in Shen (2012), following a 15-minute interaction with a robot named ‘Robovie,” children developed the idea that Robovie had feelings, interests, and was intelligent, but like objects, Robovie did not have civil liberties. Of particular relevance, results from an ERP study conducted by Chammat and colleagues (2010) suggested that emotional robot faces elicit similar ERP responses to emotionally charged human faces, even when using a non-humanoid robot. All of these results highlight how context and experience can affect the way people evaluate robots and attribute social and personality characteristics to them. Like these various behavioral results, our neural results may also be highly sensitive to the type of interactions observers have with robots, and manipulating these could offer important insights into how visual mechanisms for face processing intersect with more general mechanisms for social cognition.

Overall, our data demonstrate how the tuning of neural mechanisms for face recognition is complex. Robot faces are a useful stimulus class for examining a wide range of perceptual and social factors that affect face processing, largely because of their residence at the border between face and object. As robotic agents become more common in a range of everyday settings, understanding the nature of this boundary and what factors can affect its topography will be increasingly important.

